# Early Alpine occupation backdates westward human migration in Late Glacial Europe

**DOI:** 10.1101/2020.08.10.241430

**Authors:** Eugenio Bortolini, Luca Pagani, Gregorio Oxilia, Cosimo Posth, Federica Fontana, Federica Badino, Tina Saupe, Francesco Montinaro, Davide Margaritora, Matteo Romandini, Federico Lugli, Andrea Papini, Marco Boggioni, Nicola Perrini, Antonio Oxilia, Riccardo Aiese Cigliano, Rosa Barcelona, Davide Visentin, Nicolò Fasser, Simona Arrighi, Carla Figus, Giulia Marciani, Sara Silvestrini, Federico Bernardini, Jessica C. Menghi Sartorio, Luca Fiorenza, Jacopo Moggi Cecchi, Claudio Tuniz, Toomas Kivisild, Fernando Gianfrancesco, Marco Peresani, Christiana L. Scheib, Sahra Talamo, Maurizio D’Esposito, Stefano Benazzi

## Abstract

The end of the Last Glacial Maximum (LGM) in Europe (~16.5 ka ago) set in motion major changes in human culture and population structure^1^. In Southern Europe, Early Epigravettian material culture was replaced by Late Epigravettian art and technology about 18-17 ka ago at the beginning of southern Alpine deglaciation, although available genetic evidence from individuals who lived ~14 ka ago^2–5^ opened up questions on the impact of migrations on this cultural transition only after that date. Here we generate new genomic data from a human mandible uncovered at the Late Epigravettian site of Riparo Tagliente (Veneto, Italy), that we directly dated to 16,980-16,510 cal BP (2σ). This individual, affected by a low-prevalence dental pathology named focal osseous dysplasia, attests that the very emergence of Late Epigravettian material culture in Italy was already associated with migration and genetic replacement of the Gravettian-related ancestry. In doing so, we push back by at least 3,000 years the date of the diffusion in Southern Europe of a genetic component linked to Balkan/Anatolian refugia, previously believed to have spread during the later Bølling/Allerød warming event (~14 ka ago^4,6^). Our results suggest that demic diffusion from a genetically diverse population may have substantially contributed to cultural changes in LGM and post-LGM Southern Europe, independently from abrupt shifts to warmer and more favourable conditions.

---

During the LGM (~30-16.5 ka ago^7^) European hunter-gatherers coped with dramatic environmental and climatic upheavals by either migrating or adapting to extreme ecological conditions and reconnecting into large-scale networks^4,8,9^. Following the beginning of deglaciation large zones of the Alps were recolonised by hunter-gatherers. On the eastern Italian side this process started ~17 ka ago from the pre-Alpine valley bottoms^10,11^ and then reached altitudes as high as 1700 m above mean sea level (amsl) in the Bølling-Allerød temperate interstadial as a consequence of the progressive timberline rise^12^ (Supplementary Information section 1). In this time frame, material culture of Italy and Balkan/eastern Europe shows change over time in raw material procurement, lithic production, subsistence strategies, funerary practices, and variability of portable art, all of which document the transition from Early to Late Epigravettian^13–15^.

Current understanding of this cultural shift links the Late Epigravettian culture in northeastern Italy to a major population turnover, starting around 14ka ago at Riparo Villabruna^16^ (Vercellotti et al., 2008). It was explained as the effect of migrations from East, putatively facilitated by the Bølling-Allerød event, a warmer and more favourable period that began only about 14.7 ka ago^4,5^. Nevertheless, the processes underlying the beginning of Late Epigravettian culture and technology in the southern Alpine and Adriatic area started around 3,000 years earlier (~18-17ka ago) and are far from being disentangled due to the still fragmentary evidence for interactions between hunter-gatherers located on opposite sides of the Great Adriatic-Padanian Region^17^ (in Italy and the Balkans).

To understand the full extent of the role played by demic processes in this key transition in Late Glacial Europe we focused on the left hemimandible of an individual found at Riparo Tagliente (Tagliente2^18^) associated with Late Epigravettian evidence (Supplementary Information section 2). During the LGM and the Late Glacial, the Adriatic Sea basin played a critical role in shaping the economy and mobility of Epigravettian groups. Geomorphological and sedimentological processes linked to the extension of Alpine glaciers and to the lowering of the sea level at ~120m amsl formed a vast plain area that connected the Italian and Balkan peninsulas. The central Slovenian passage was used by large pachyderms and ruminants such as cervids and bisons which, alongside with bears, were the main game targeted by Epigravettian hunter-gatherers^19^. Riparo Tagliente (northeastern Italy) represents the earliest available evidence of human occupation of the southern Alpine slope^11^ while the main glaciers in the area started withdrawing 17.7 ka ago^20^, and is therefore critical to address questions on the impact of human movement in this time frame (Fig.1). We performed anthropological and genetic analyses to assess the biological background of the sampled individual. The hemimandible was also directly dated to independently ascertain its chronology and the possible contemporaneity with contextual post-cranial human remains from a partially preserved burial (Tagliente1^21^; Supplementary Information section 2).

**Fig. 1.**
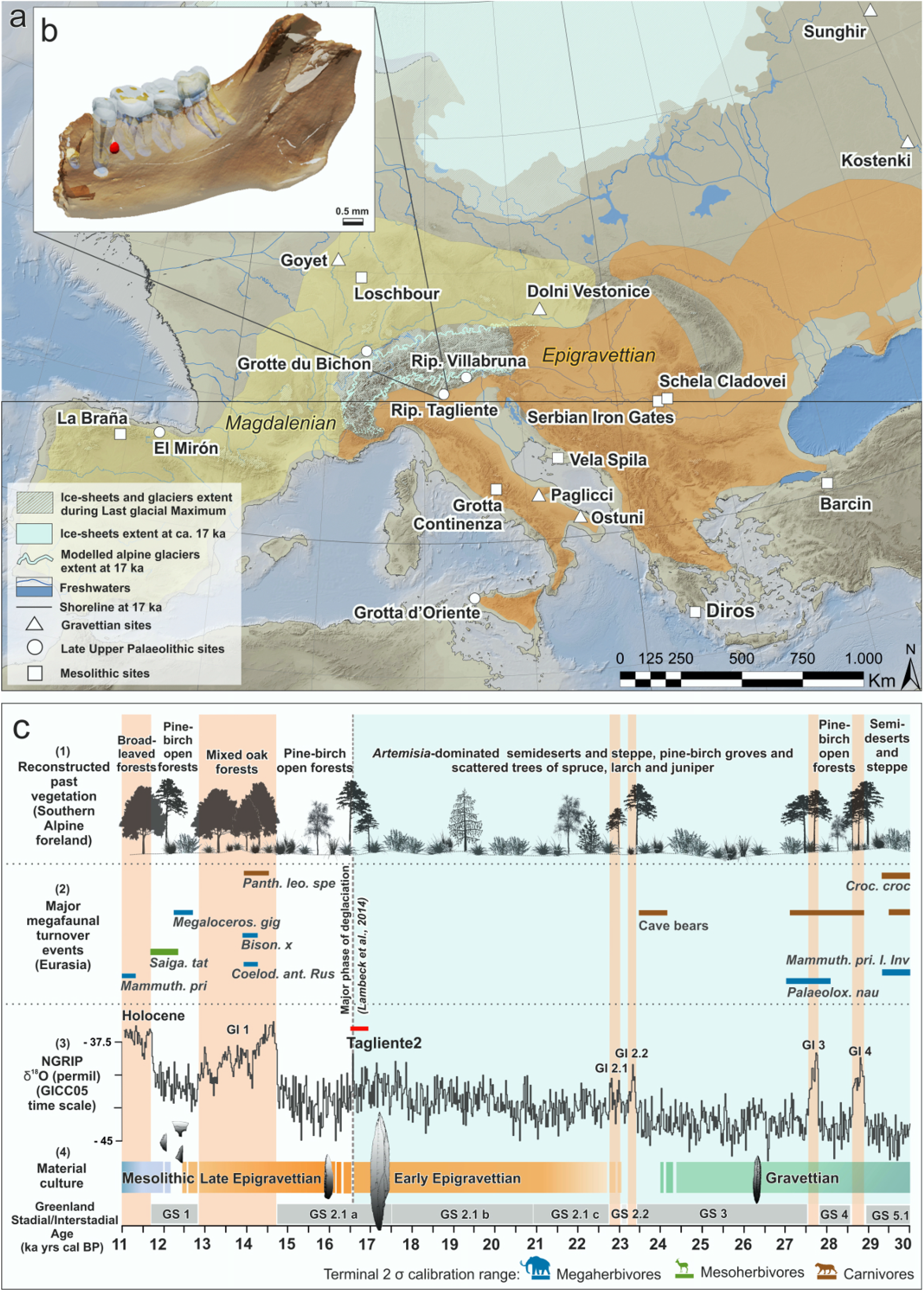
Geographical, ecological, and cultural context of the study. a) Palaeogeographic map of Europe during Late Glacial, centered at 17 ka ago. Coordinate system ETRS89 / UTM zone 32N (EPSG 25832); Digital Elevation Model (base topography – Copernicus Land Monitoring Service 2019 (CLMS, http://land.copernicus.eu/paneuropean), and General Bathymetric Chart of the Oceans (GEBCO 2019 grid, doi:10.5285/836f016a-33be-6ddc-e053-6c86abc0788e). Sea level drop at – 110 m^7^. Scandinavian and British Islands ice sheets, mountain glaciers Last Glacial Maximum (LGM) extent (striped areas) and freshwater systems modified after^22^. Scandinavian and British Islands ice sheets (pale blue) at 17 ka after^23^. Alpine glaciers extent (dashed outline) modelled at 17 ka from^24^ (https://doi.org/10.5446/35164). Modelled extension is generally underestimated in the northwestern Alps and overestimated in the eastern and south-western Alps^24^. Coloured areas refer to the distribution of Epigravettian and Magdalenian material culture at 17ka ago, while white symbols indicate the geographic location of the main sampling sites discussed in the text, encompassing a period ranging from ~30 to 8ka. b) 3D lateral view of the hemimandible Tagliente2. Reduced opacity shows roots, pulp chambers, dentine and enamel of the preserved teeth, as well as the cementome in red between the distal side of P_3_ root and the mesial root side of M_1_. c) Comparison between palaeoclimate, palaeoenvironmental proxies, and cultural proxies over the 30-11 ka cal BP time span. Key to panels: (1) Reconstruction of past vegetation based on Southern Alpine foreland palaeoecological records^12,22,25,26^; (2) Eurasian major megafaunal transitions (regionwide extirpations or global extinctions, or invasions, of species or major clades) identified in Late Pleistocene Holarctic megafaunal data sets through a DNA or paleontological studies^19,27^; (3) NGRIP δ^18^O record in 20 yrs means on the GICC05 time scale^28^; (4) Material cultural sequence for Eastern and Southern Europe. The chronological interval obtained for Tagliente2 is indicated by a red rectangle (16,980-16,510yrs cal BP, 2σ interval, IntCal20^29^).

## Tagliente2

The left hemimandible Tagliente2 is mesially broken to the alveolus of the first premolar (P_3_), while the ramus is mostly complete, except for the condyle and coronoid process. Four permanent teeth (P_4_-M_3_) are still in place into their respective alveoli, but only a tiny apical portion of the LP_3_ root is preserved. The presence of wear pattern on the third molar (wear stage 2^30^) and its eruption suggest that the mandible can be ascribed to a young adult, while the robusticity index (42.59, this study) and the gonial angle (110°) fall within the range of male variability^18^. X-ray microCT scans and digital segmentation of Tagliente2 (Extended Data Fig.1) identified the presence of a rounded alteration close to the buccodistal aspect of the P_4_ root. The identified lesion shows homogeneous radiopacity surrounded by a radiotransparent thin area (Extended Data Fig.2) and consists of an irregular (Volume = 10.71 mm^3^; bucco-lingual diameter= 2.48cm; mesio-distal diameter = 2.90cm; see Extended Data Fig.1) and compact dental tissue which was mechanically removed (see Methods) and physically analysed through histological examination (see Methods). The latter suggests the presence of stratified and acellular cementum material (Extended Data Fig.2), which together with anatomical position and morphology (Fig.1b) provides evidence ascribable to focal cemento-osseous dysplasia (FCOD; Supplementary Information section 3), a benign lesion of the bone in which normal bone is replaced by fibrous tissue, followed by calcification with osseous and cementum tissue^31^ (Supplementary Information section 4).

## Radiocarbon Dating

The hemimandible of Tagliente2 was directly dated to 16,980-16,510 cal BP (95.4% probability using IntCal20^29^ in OxCal v.4.3^32^) confirming the attribution to the Late Epigravettian chronological range. This result is consistent with the associated material culture, and with the attribution to the same cultural context as Tagliente1 (16,130–15,560 cal BP^21^; Supplementary Information section 5).

## Ancient DNA

We extracted DNA from five samples taken from mandibular and tooth tissues and screened for the presence of endogenous human DNA through a pooled whole genome sequencing. One of the healthy mandibular samples yielded sufficient endogenous DNA (5.06%) and was re-sequenced to achieve a total genome wide coverage of 0.28x, yielding 266K SNPs overlapping with the *Human Origins* SNP Array. We also provide a number of non-reference sites found to overlap genes known to be involved with cementoma insurgence, which, given the low coverage and ancient DNA degradation, are reported here with no further interpretation.

Overall contamination estimated from mtDNA was 2.158% and 0.60 – 1.53% from the X chromosome. The mtDNA haplogroup is a basal U4’9, consistent with a European Palaeolithic individual (Fig. 2A). X/Autosome coverage ratio in the order of 0.56 confirmed the individual was male, and the chromosome Y haplogroup estimated to be I2, which captures the majority of the post-Villabruna diversity in Europe (Fig. 2B). From a population perspective, we performed a MDS analysis based on outgroup f3 distances (Fig. 3A) and found the sample to fall within the broader European Western Hunter Gatherer (WHG) genetic variation, pointing to an affinity to the previously described Villabruna Cluster^4^, known to have largely replaced previous European Hunter Gatherer populations at least ~14 ky ago. One of the defining feature of the Villabruna cluster is a higher affinity with Near Eastern genetic components, compared with pre-existing palaeolithic West Eurasians: the significantly negative f4 test (Kostenki14, Tagliente2, Druze, Mbuti: f4=−0.0037; standard error= 0.00063; Z=−5.88) further confirmed Tagliente2 to share genetic features with the Villabruna cluster and to be in discontinuity with the preceding European genetic background. We followed up this observation using a series of f4 tests in the form (Tagliente2, X; Y, Mbuti), where Y is a population of interest and X is either Villabruna (~14ky ago^4^), Bichon (~13.7ky ago^4^) or a Mesolithic Italian from Grotta Continenza (~11.9ky ago^33^; Fig. 3 B). We chose three independent WHG samples to control for potential biases introduced by the genotyping strategy, and indeed found small discrepancies when we compared results obtained using capture (Villabruna) or shotgun (Bichon and Continenza) data. To minimise this effect we chose to put more weight on the interpretation of shotgun results, deemed to be more readily comparable with the shotgun data generated for Tagliente2 in this study. We also show that the data available is sufficient to achieve significance in a f4 test when the order of X and Y populations are inverted, as for (Tagliente, Y, Grotta Continenza, Mbuti, Extended Data Fig.3). The higher affinity of Continenza and Bichon to later WHG (Loschbour, Iberia_HG and Continenza, Bichon, and Villabruna themselves) when compared to Tagliente2 may be explained by the more ancient age of Tagliente2 or with the former individuals being genetically closer to the genetic ancestry that reached central Europe at least by 14ka ago. Alternatively, the higher affinity emerging among more recent WHG samples may also be ascribed to subsequent admixtures between the newly arrived Tagliente2 individuals and pre-existing Dolní Věstonice- or Goyet-like (~35ka ago^4^) genetic substrates, as already reported for Loschbour (~8.1ka ago^4,34^). We then modelled the position of Tagliente2 within the tree proposed by Fu and colleagues (2016; Extended Data Fig.4A) and found that it may fit well within the Villabruna branch confirming previous results (Extended Data Fig.4B). Notably, alternative qpGraphs in which Villabruna was modelled as a mixture between Tagliente2 and Věstonice (Extended Data Fig.4C) or between Tagliente2 and Goyet (Extended Data Fig.4D) yield comparable summary statistics and would allow for up to 10% contribution from Věstonice or up to 5% contribution from Goyet into the later samples belonging to the Villabruna cluster. To minimise the effects of the mismatch between capture and shotgun data due to attraction, we also explore the feasibility of the basic qpGraph (Extended Data Fig.4B) using, where possible, shotgun samples (Extended Data Fig.5).

**Fig. 2.**
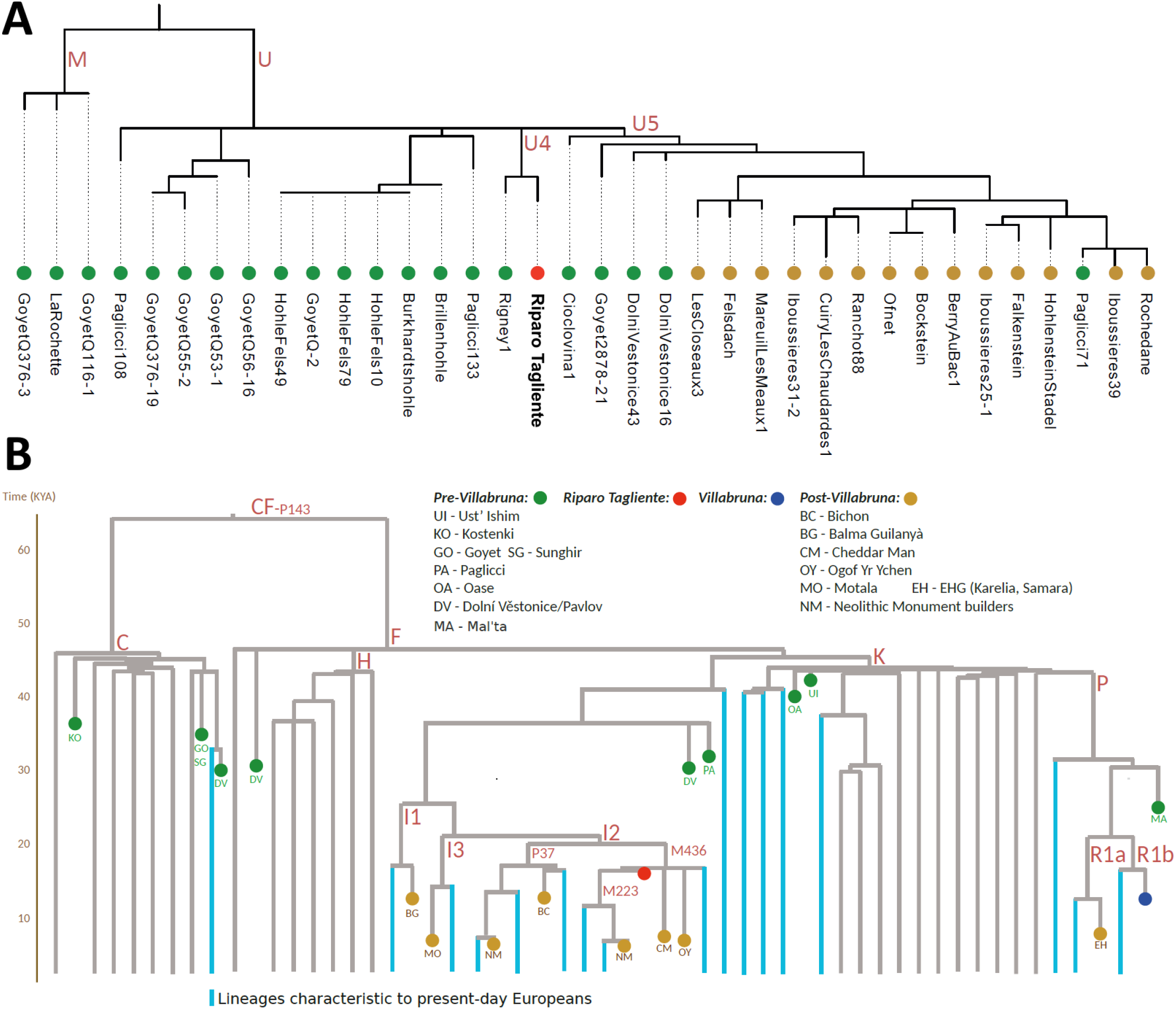
Uniparental haplogroups of Tagliente2. Panel A shows the mtDNA haplogroup of Tagliente2 (in red) within a number of pre (Green) and post (Gold) Villabruna samples. Panel B shows the chrY haplogroup of Tagliente2 (in red) surrounded by post-Villabruna samples (including Bichon, BC). Y Haplogroup splits are drawn according to the dater estimates based on high coverage modern sequences from Karmin et al. 2015. The ancient individuals are mapped on this tree considering the available haplogroup-informative available SNP data and private mutations in the ancient samples have been ignored.

**Fig. 3:**
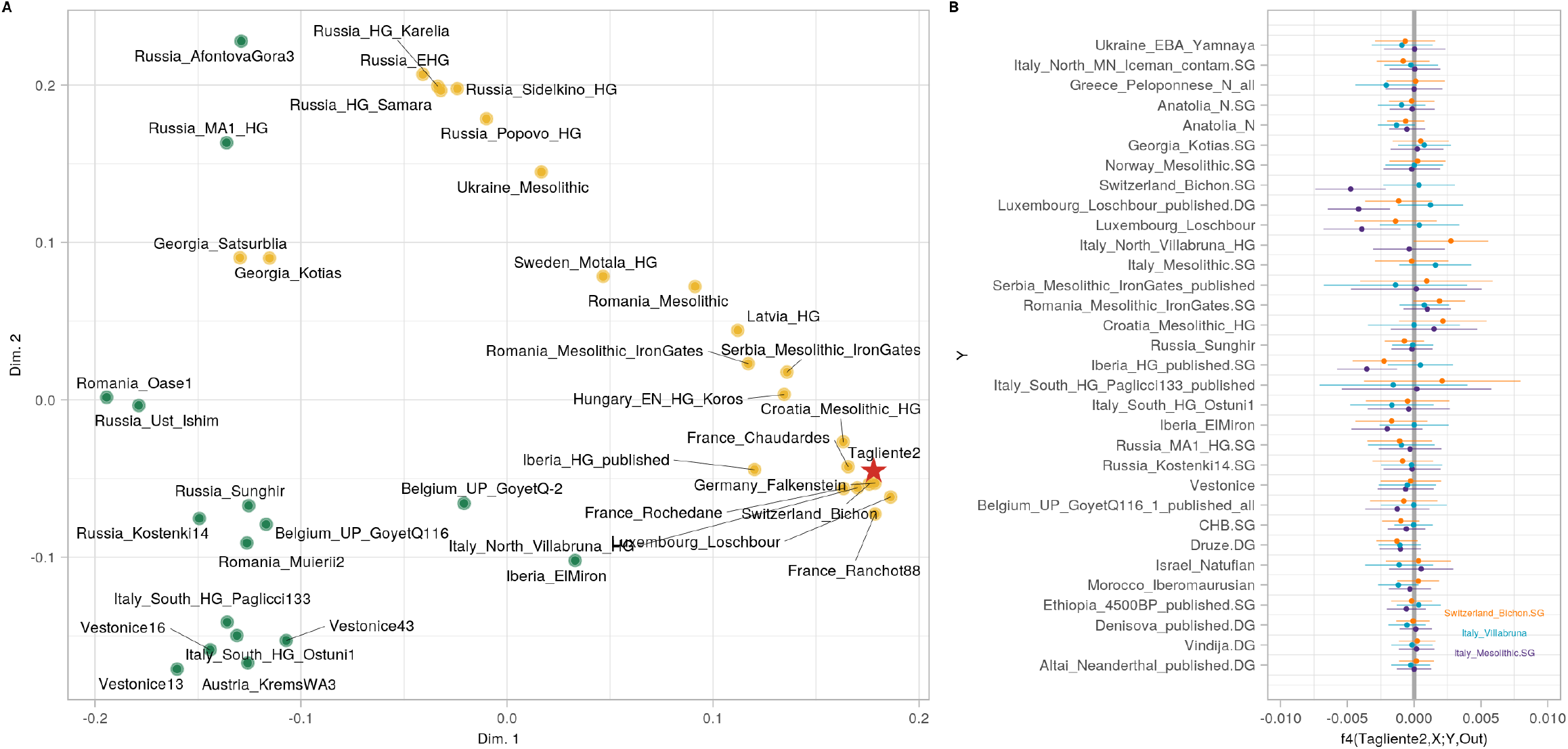
Demographic inference from Tagliente 2. A) Mutidimensional Scaling based on (Mbuti; X, Tagliente2) outgroup f3 statistics show Tagliente2 (red star) to cluster within the Villabruna Cluster (in gold) and away from the pre-existing South European samples (in green). B) f4 tests (Tagliente2, X, Y, Mbuti +/− 3 s.e.) where X is either a Mesolithic Italian, Villabruna or Bichon WHG sample and Y, shown along the y axis, is a population of interest.

## Discussion

Tagliente2 is clearly associated with Late Epigravettian material culture, and its direct date (~17ka ago) is consistent with the chronology of the lowermost layers attested at Riparo Tagliente^11^. Therefore, this individual belongs to one of the earliest human groups that first occupied the southern Alpine slopes at the end of the LGM, when mountain ranges in Southern Europe became accessible to the expansion of animals and predators^10,17^. In the same temporal interval there is evidence of a shift from Solutrean to Magdalenian material culture in Southwestern Europe, and from Early to Late Epigravettian material culture in a vast area ranging from the Rhone river to the Southern Russian plain^35^. In this broad context, the individual found at Riparo Tagliente denotes the presence in the region of human groups from Eastern Europe/Anatolia, therefore backdating by at least 3ka the occurrence of genetic components previously reported for the later Villabruna cluster (~14ka ago^2–6^). At the same time, even earlier migrations into Southern Europe might be envisaged to explain the presence of Villabruna- or Tagliente2-related genetic components in the ~18.7 ka-old sample found at El Mirón, Spain^4,34^. Tagliente2 seems to be basal to all samples that are more recent than Villabruna based on both autosomic DNA and Y chromosome. From an autosome perspective, this may result from an admixture between Villabruna and survivors of the Věstonice or Goyet clusters between 17 and 14 ka ago. Nevertheless, the symmetrical relationship of Tagliente2 and other post 14kya European hunter-gatherers with Věstonice, Ostuni, Goyet or Kostenki (Fig. 3B) does not seem to support this scenario in spite of the promising qpGraph (Extended Data Fig.4). In addition, we report slight f4 discrepancies in the analysis when the same individuals is obtained either through capture or shotgun (e.g. Anatolia_N in Fig. 3 B), or when using Bichon or Continenza versus Villabruna, all of which show the importance of avoiding capture and shotgun samples within the same f4 test.

Tagliente2 therefore provides evidence that the major migrations which strongly affected the genetic background of all Europeans^2–6^, started considerably earlier in Southern Europe than previously reported, and in this region they do not seem to be limited to favourable, warmer periods (e.g. Greenland Interstadial 1, ~14.7-13ka ago). Our results rather show that population movements were already in place during the cold phase immediately following the LGM peak. At this stage, Italy, the Balkans, and Eastern Europe/Western Asia were already connected into the same network of potential LGM refugia, and exchanged both genes and cultural information. This finding also backdates previous conclusions concerning a plausible demic component to change over time in the coeval material culture of Southern Europe^2–6^, and temporally locates this process at the transition between Early and Late Epigravettian or even possibly at an early stage of the Epigravettian and at the very beginning of the Magdalenian sequence. The latter scenario stems from interpreting the traces of Villabruna/Tagliente2 genetic components recorded at El Mirón (~19ka ago^4,34^) as the result of an early phase of this westward expansion, rather than as a proliferation from Western European refugia. Further genetic evidence from Southern European contexts dated between ~24-19 ka ago, however, will be needed in the future to test this hypothesis.

Discontinuity between Early Epigravettian and Late Epigravettian lithic tool manufacturing is documented since ~18-17ka cal BP in Southern and Eastern Europe, despite the biased spatial and temporal distribution of the archaeological record^13^. In the period of interest, however, human groups inhabiting Southern Europe were exposed to limited ecological risk^8,36^ and in this time frame there is at present a lack of evidence of their direct impact on megafaunal extinction, which is consistently associated with short interstadial (warming) events during the LGM and following the beginning of the Bølling/Allerød event^19,27,28^. In this context, the emergence of Late Epigravettian lithic technology may have been less prone to local adaptation than to the transmission of culture via population movement and human interaction^37^. Tagliente2, therefore, suggests that cumulative cultural change observed in Southern Europe from the end of LGM to the end of the Younger Dryas (~11.7 ka ago) was at least in part triggered by gene flow from eastern refugia into Northeastern Italy and that this process, in its early stage, was independent of warming events, and contributed to the gradual replacement of pre-LGM ancestry across the Italian peninsula^38^.

Furthermore, our analysis offers new insights on the presence and distribution of a rare dental pathology (FCOD) that has never been reported before in human fossils and is today more frequently associated with female individuals of African American or Southeastern Asian descent^39^. Although etiology and developmental trajectory of this pathologic condition are yet to be understood, this work documents its presence in pre-Neolithic and pre-industrial societies of Europe.

In conclusion, the present results open a debate on the impact of demic diffusion on the very origin of the Late Epigravettian material culture and the mechanisms underlying patterns recorded before and after the onset of the Bølling/Allerød event. These include the later convergent emergence across Eurasia of a more flexible and responsive technology, and of schematic art^40^ associated with change in demographic pressure, environmental challenges, and mobility^41–43^.

## Methods

### Radiocarbon dating

The teeth from the hemimandible (Tagliente2) were pretreated at the Department of Human Evolution at the Max Planck Institute for Evolutionary Anthropology (MPI-EVA), Leipzig, Germany, using the method previously published^44^. Circa 500 mg of the whole root of the tooth is taken. The sample is then decalcified in 0.5M HCl at room temperature until no CO_2_ effervescence is observed. 0.1M NaOH is added for 30 minutes to remove humics. The NaOH step is followed by a final 0.5M HCl step for 15 minutes. The resulting solid is gelatinized following Longin (1971)^45^ at pH3 in a heater block at 75°C for 20h. The gelatine is then filtered in an Eeze-FilterTM (Elkay Laboratory Products (UK) Ltd.) to remove small (<80 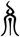 m) particles. The gelatine is then ultrafiltered^46^ with Sartorius “VivaspinTurbo” 30 KDa ultrafilters. Prior to use, the filter is cleaned to remove carbon containing humectants^47,48^. The samples are lyophilized for 48 hours. The date is corrected for a residual preparation background estimated from ^14^C free bone samples. These bones were kindly provided by D. Döppes (MAMS, Germany), and one was extracted along with the batch from the tooth^49^. To assess the preservation of the collagen yield, C:N ratios, together with isotopic values must be evaluated. The C:N ratio should be between 2.9 and 3.6 and the collagen yield not less than 1% of the weight^50^. For the tooth stable isotopic analysis is evaluated at MPI-EVA, Leipzig (Lab Code R-EVA 1606) using a ThermoFinnigan Flash EA coupled to a Delta V isotope ratio mass spectrometer. The Tagliente tooth passed the collagen evaluation criteria and between 3 and 5 mg of collagen inserted into pre-cleaned tin capsules. This was sent to the Mannheim AMS laboratory (Lab Code MAMS) where was graphitized and dated^51^.

### DNA Extraction and Sequencing

The DNA extraction and sample library was prepared in the dedicated ancient DNA laboratory at the Estonian Biocentre, Institute of Genomics, University of Tartu, Tartu, Estonia. The library quantification and sequencing were performed at the Estonian Biocentre Core Laboratory. The main steps of the laboratory work are detailed below.

### DNA extraction

Tooth/bone material was powdered at the aDNA clean lab of the Department of Cultural Heritage, University of Bologna by G.O. and S.S. and sent to the University of Tartu. To approximately 20 mg of powder 1000 μl of 0.5M EDTA pH 8.0 and 25 μl of Proteinase K (18mg/ml) were added inside a class IIB hood. The sample was incubated for 24 h on a slow shaker at room temperature. DNA extracts were concentrated to 250 μl using Vivaspin® Turbo 15 (Sartorius) concentrators and purified in large volume columns (High Pure Viral Nucleic Acid Large Volume Kit, Roche) using 10X (2.5 ml) of PB buffer (Qiagen) following the manufacturers’ instructions with the only change being a 10 minute incubation at 37 degrees prior to the final elution spin and eluted in 100 μl of EB buffer (QIAGEN). Samples were stored at −20 C.

### Library preparation

The extracts were built into double-stranded, single-indexed libraries using the NEBNext® DNA Library Prep Master Mix Set for 454™ (E6070, New England Biolabs) and Illumina-specific adaptors52 following established protocols^52–54^. DNA was not fragmented and reactions were scaled to half volume, adaptors were made as described in^52^ and used in a final concentration of 2.5uM each. DNA was purified on MinElute columns (Qiagen). Libraries were amplified using the following PCR set up: 50μl DNA library, 1X PCR buffer, 2.5mM MgCl2, 1 mg/ml BSA, 0.2μM inPE1.0, 0.2mM dNTP each, 0.1U/μl HGS Taq Diamond and 0.2μM indexing primer. Cycling conditions were: 5’ at 94C, followed by 18 cycles of 30 seconds each at 94C, 60C, and 68C, with a final extension of 7 minutes at 72C. Amplified products were purified using MinElute columns and eluted in 35 μl EB (Qiagen). Three verification steps were implemented to make sure library preparation was successful and to measure the concentration of dsDNA/sequencing libraries – fluorometric quantitation (Qubit, Thermo Fisher Scientific), parallel capillary electrophoresis (Fragment Analyser, Advanced Analytical) and qPCR.

Library quality and quantity have been assessed by using Agilent Bioanalyzer 2100 High Sensitivity and Qubit DNA High Sensitivity (Invitrogen). The initial shotgun screening was done on NextSeq500 using the High-Output 75 cycle single-end kit. The secondary, paired-end sequencing was performed on the NovaSeq6000 (Illumina), flowcell S1, without any other samples to ensure no index-hopping due to the single-indexing of the sample, generating 150-bases paired-end reads. Whole genome sequencing with Illumina paired-end (2×150 bp) led to 488 million high-quality reads. About 6.99% of the reads could be successfully mapped on the human genome sequence with a duplication rate of 51%, leading to an average 0.28x genome coverage. The mapped reads showed nucleotide misincorporation patterns which were indicative of post-mortem damage (Extended Data).

### Sequencing filtering, mapping and variant calling analysis

Before mapping, the paired end reads were merged and corrected using FLASH^55^. The merged reads were trimmed of adapters, indexes and poly-G tales occuring due to the specifics of the NextSeq500 and NovaSeq technology using cutadapt-1.11^56^. Sequences shorter than 30 bp were also removed with the same program to avoid random mapping of sequences from other species. The sequences were aligned to the reference sequence GRCh37 (hs37d5) using Burrows-Wheeler Aligner (BWA 0.7.12)^57^ and the command mem with seeding disabled. After alignment, the sequences were converted to BAM format and only sequences that mapped to the human genome were kept with samtools-1.3^58^. Afterwards, the data from different flow cell lanes were merged and duplicates were removed using picard 2.12 (http://broadinstitute.github.io/picard/index.html). Indels were realigned using GATK-3.5^59^ and reads with a mapping quality less than 10 were filtered out using samtools-1.3 (ref). In order to maximise the coverage of sites included in the 1240 Human Origin capture array, a random read with mapping quality above 30 and phred score above 33 was chosen to represent the pseudo-haploid genotype of our sample, and then merged with reference data from ancient and modern European samples.

### aDNA authentication

As a result of degrading over time, aDNA can be distinguished from modern DNA by certain characteristics: short fragments and a high frequency of C=>T substitutions at the 5’ ends of sequences due to cytosine deamination. The program mapDamage2.0^60^ was used to estimate the frequency of 5’ C=>T transitions. Rates of contamination were estimated on mitochondrial DNA by calculating the percentage of non-consensus bases at haplogroup-defining positions as detailed in (ref). Each sample was mapped against the RSRS downloaded from phylotree.org and checked against haplogroup-defining sites for the sample-specific haplogroup.

Samtools 1.9^58^ option stats was used to determine the number of final reads, average read length, average coverage etc.

### Calculating genetic sex estimation

Genetic sex was calculated using the methods described in^61^,estimating the fraction of reads mapping to Y chromosome out of all reads mapping to either X or Y chromosome. Additionally, sex was determined using a method described in^62^, calculating the X and Y ratio by the division of the coverage by the autosomal coverage.

### Determining mtDNA and Y chromosome haplogroups

Mitochondrial DNA haplogroups were determined using Haplogrep2 on the command line. For the determination, the reads were re-aligned to the reference sequence RSRS and the parameter --rsrs were given to estimate the haplogroups using Haplogrep2 [83,84]. Subsequently, the identical results between the individuals were checked visually by aligning mapped reads to the reference sequence using samtools 0.1.19^58^ command tview and confirming the haplogroup assignment in PhyloTree.

A total of 5703 Y chromosome haplogroup informative variants^63,64^ from regions that uniquely map to Y chromosome were covered by at least one read in the sample and these were called as haploid from the BAM file using the --doHaploCall function in ANGSD^65^. Derived and ancestral allele and haplogroup annotations for each of the called variants were added using BEDTools 2.19.0^66^ intersect option. Haplogroup assignments of each individual sample were made by determining the haplogroup with the highest proportion of informative positions called in the derived state in the given sample.

**MDS, f3, f4 and qpGraph** Reference samples were downloaded from https://reich.hms.harvard.edu/downloadable-genotypes-present-day-and-ancient-dna-data-compiled-published-papers and merged with the newly generated Tagliente2 data. A set of Outgroup f3^67^ in the form (X, Y, Mbuti) was run on samples listed in Fig.3A, and the resulting pairwise distance matrix (distance=1-f3) was used to compute a Multi Dimensional Scaling (MDS). The f4 test^67^ was run using the popstats.py script^68^. qpGraph were generated using Admixtools^67^, starting from the backbone described in Fu et al. 2016 and investigating putative positions for Tagliente2 as informed by the f4 results described in Fig. 3B.

### Histopathological examination

A thin section of 0.005 mm was sampled for histological analysis. The cross-section has been performed following mesiodistal direction based on the best location for core sampling. The core was fixed in buffered neutral formalin 10% in order to protect the fibrous elements of cementum from damage caused by the acids used as decalcifying agent performed with Trichloroacetic acid for 7 days. Finally, the section was coloured by Hematoxylin / Eosin. The sample was mounted on frosted glass slides and thin-sectioned using a Struers Accutom-50. The preparation of the histological section was carried out at the Centro Odontoiatria e Stomatologia F. Perrini, Pistoia, Italy. the section was studied using a Nikon E200 microscope. Photomicrographs were captured using NIS D 3.0 Software and edited in Adobe Photoshop CC.

### Data Availability

Whole genome sequences generated for this study are freely available for download at www.xxx.xx/xxxx_xxxx/xxxx_xxx/

## Acknowledgements

This work was supported by the European Union through the European Research Council under the European Union’s Horizon 2020 Research and Innovation Programme (grant agreement No 724046 – SUCCESS awarded to S.B. - www.erc-success.eu; grant agreement No. 803147 RESOLUTION awarded to S.T., https://site.unibo.it/resolution-erc/en) as well as through the European Regional Development Fund (Project No. 2014–2020.4.01.16–0030 to C.L.S. and T.S.) and projects No. 2014-2020.4.01.16-0024, MOBTT53 (L.P.); by the Estonian Research Council personal research grant [PRG243] (C.L.S); by UniPd PRID 2019 (L.P.). We thank the Ministry of Cultural Heritage and Activities of Italy, and Veneto Archaeological Superintendency for granting access to the human remains of Tagliente2, in particular the SABAP of Verona, Rovigo and Vicenza. We are grateful to Dr. Francesca Rossi and Dr Nicoletta Martinelli for giving access to human remains from Riparo Tagliente and for the support they provided during sampling. We also thank Dr. Elisabetta Cilli and Andrea De Giovanni for assistance with the preliminary documentation of the hemimandible Tagliente2.

## Author contribution

Designed Study: E.B., L.P., G.O., S.B., Run Analyses: E.B., L.P., G.O., C.P., T.S., F.M., T.K., C.S., S.T., Wrote Manuscript: E.B., L.P., G.O., F.F., F.Ba., M.R., F.L., M.P., S.B., Provided Samples, Reagents or Sequences: F.F., R.A., C.S., S.T., M.D., S.B., Contributed Interpretation of Results: C.P., F.F., F.Ba., D.M., M.R., F.L., A.P., M.B., N.P., A.O., S.A., C.F., G.M., S.S., F.Be., J.Me., L.F., J.Mo., C.T., T.K., F.G., M.P., M.D.

## Competing interest declaration

The authors declare no competing interests

---

**Supplementary Information** is available for this paper

**Extended Data Fig.1.**
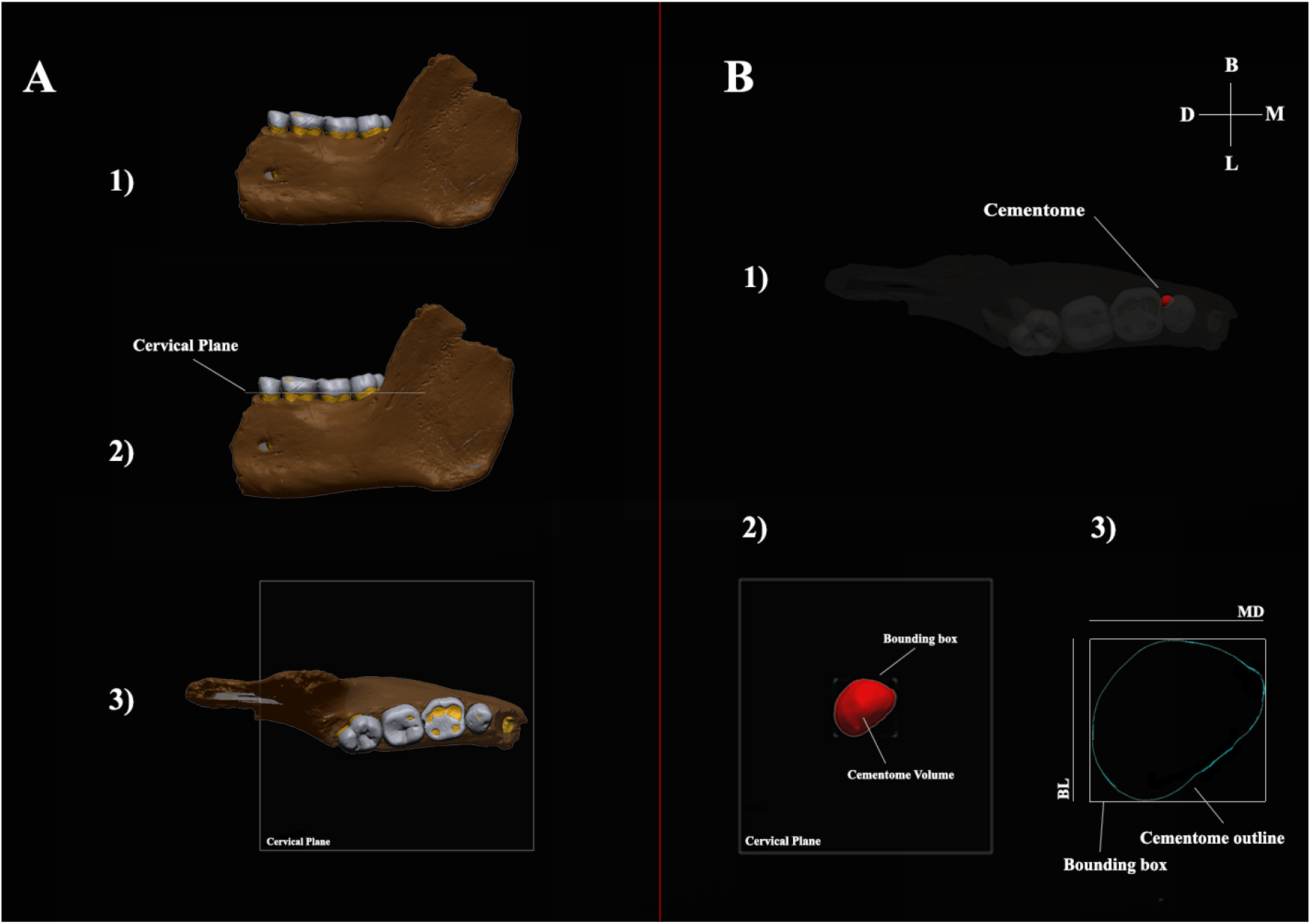
Digital reconstruction of the hemi-mandible. **A1)** A spline curve was digitized at the cervical line of each crown dentine. **A2)** A best-fit plane (cervical plane) was obtained. **A3)** The hemimandible was oriented with the best-fit plane computed at the cervical lines (i.e., the cervical plane that best fits a spline curve digitized at the cervical line), parallel to the xy-plane of the Cartesian coordinate system and rotated along the z-axis to have its lingual aspect parallel to the x-axis. **B1)** Virtual model oriented in occlusal view. Transparency level set at 72% to highlight the lesion (in red). **B2)** Cementome in occlusal view. It was excluded from the context to calculate the Volume and diameters. **B3)** the outline corresponds to the silhouette of the oriented cementome as seen in occlusal view and projected onto the cervical plane. The contour of the section identified by the cervical plane represents the cementome outline. The size of the bounding box enclosing the silhouette was used to collect mesiodistal (MD) and buccolingual (BL) diameters. M= Mesial; B= Buccal; = D=Distal; L=Lingual.

**Extended Data Fig.2.**
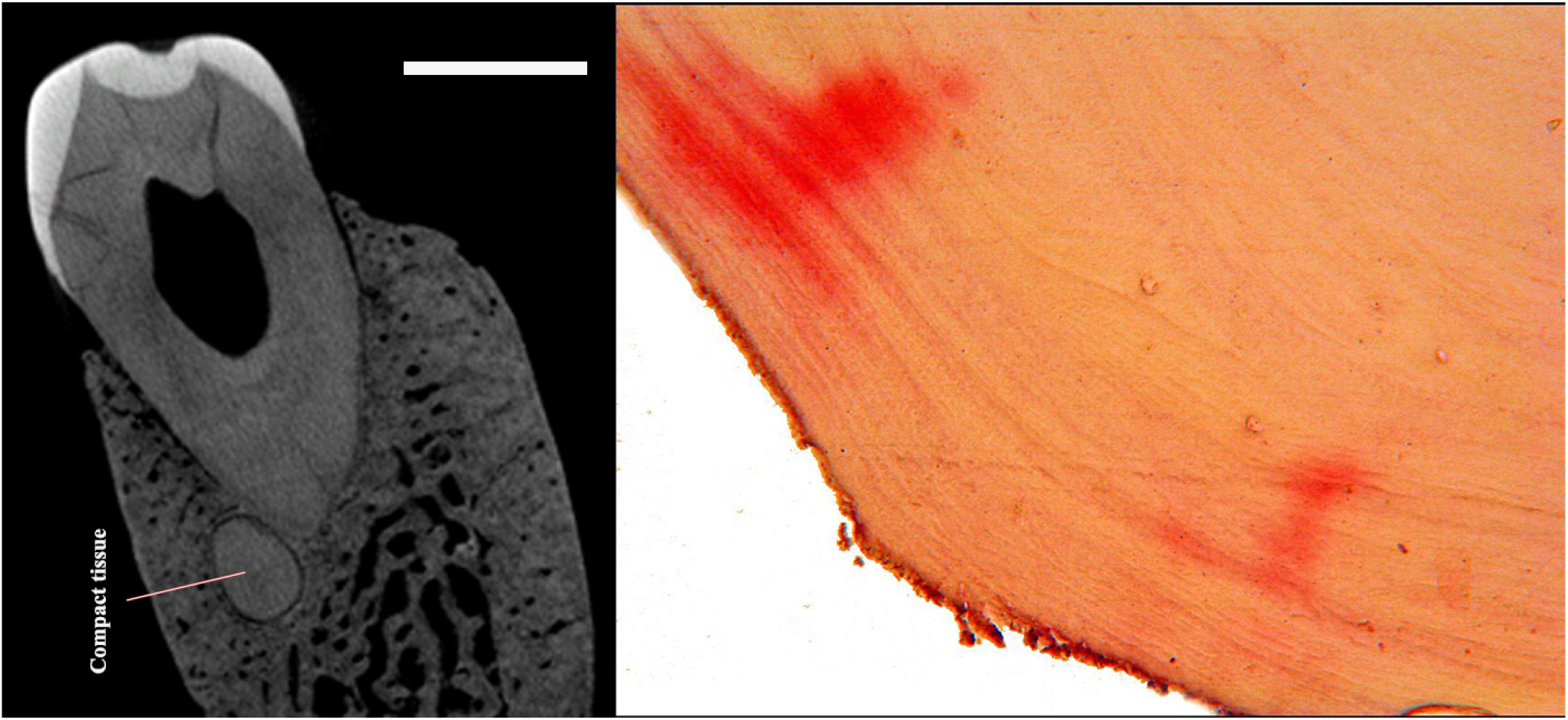
Tagliente 2 virtual (left) and physical (right) section. On the left, microCT distal view of the premolar and its pathological cementum tissue. On the right, histological section. Magnification (250X) of the cementum tissue colored by Hematoxylin/Eosin. B = Buccal; L = Lingual; scalebar = 0.5mm.

**Extended Data Fig.3:**
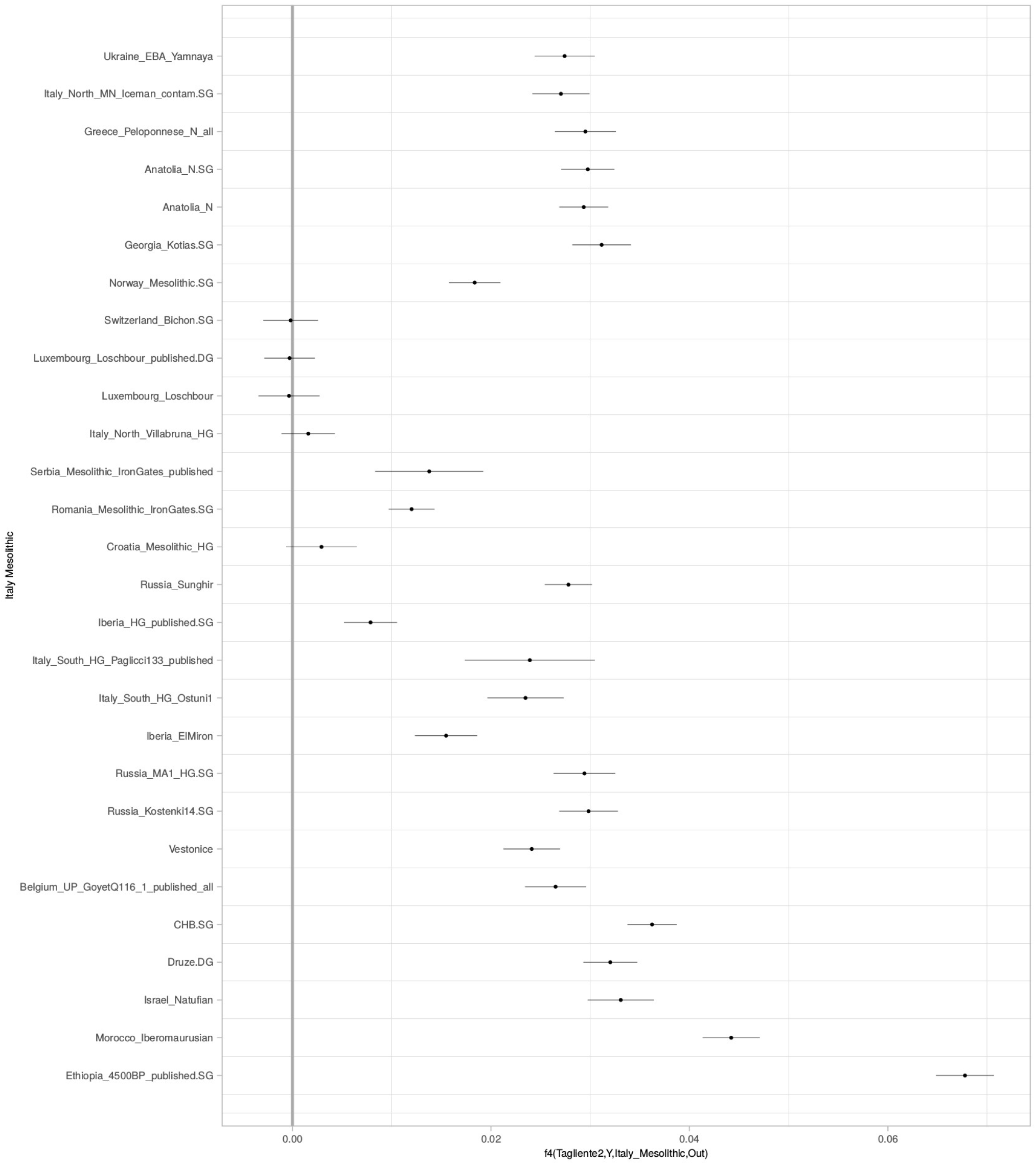
F4 tests in form (Tagliente, X, Mesolitic_Italian_Continenza, Mbuti), where X is one of the populations reported along the Y axis and consistent with Figure 3 B, showing that the available SNPs are sufficient to yield significance in a f4 test.

**Extended Data Fig.4:**
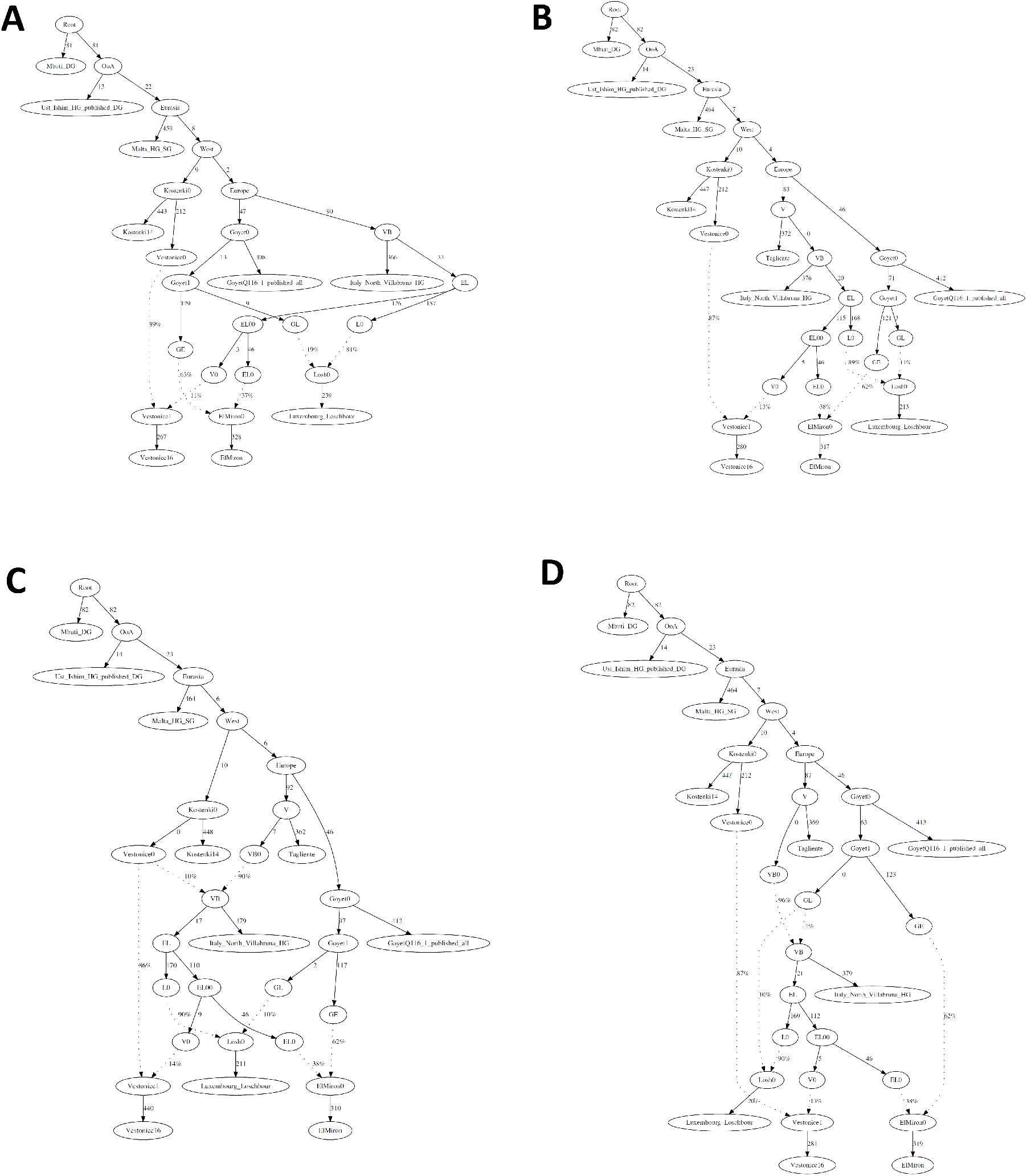
Tagliente2 within the Fu et al. 2016 qpGraph model. We started from the qpGraph originally proposed in Fu et al. 2016 (Panel A: Final Score 37628.736; dof: 8; no f2 outliers; worst f4: Mbu,Ust,Goy,Ita Z=−3.330) and informed by the f4 stats shown in Figure 3 placed Tagliente 2 as a basal branch of the Villabruna cluster (Panel B: Final Score 38182.359; dof: 15; no f2 outliers; worst f4: Mbu,Ust,Goy,Ita Z=−3.330). We also explored alternative scenarios featuring Villabruna as an admixture of pre-existing Vestonice (Panel C: Final Score 39471.021; dof: 14; no f2 outliers; worst f4: Mbu,Ust,Goy,Ita Z=−3.330) or Goyet (Panel D: Final Score 37245.412; dof: 14; no f2 outliers; worst f4: Ust,Ita,Goy,Ita Z= 3.503) clusters. Notably, since the number of events and degrees of freedom (dof) are different across different graphs, final scores are not directly comparable.

**Extended Data Fig.5:**
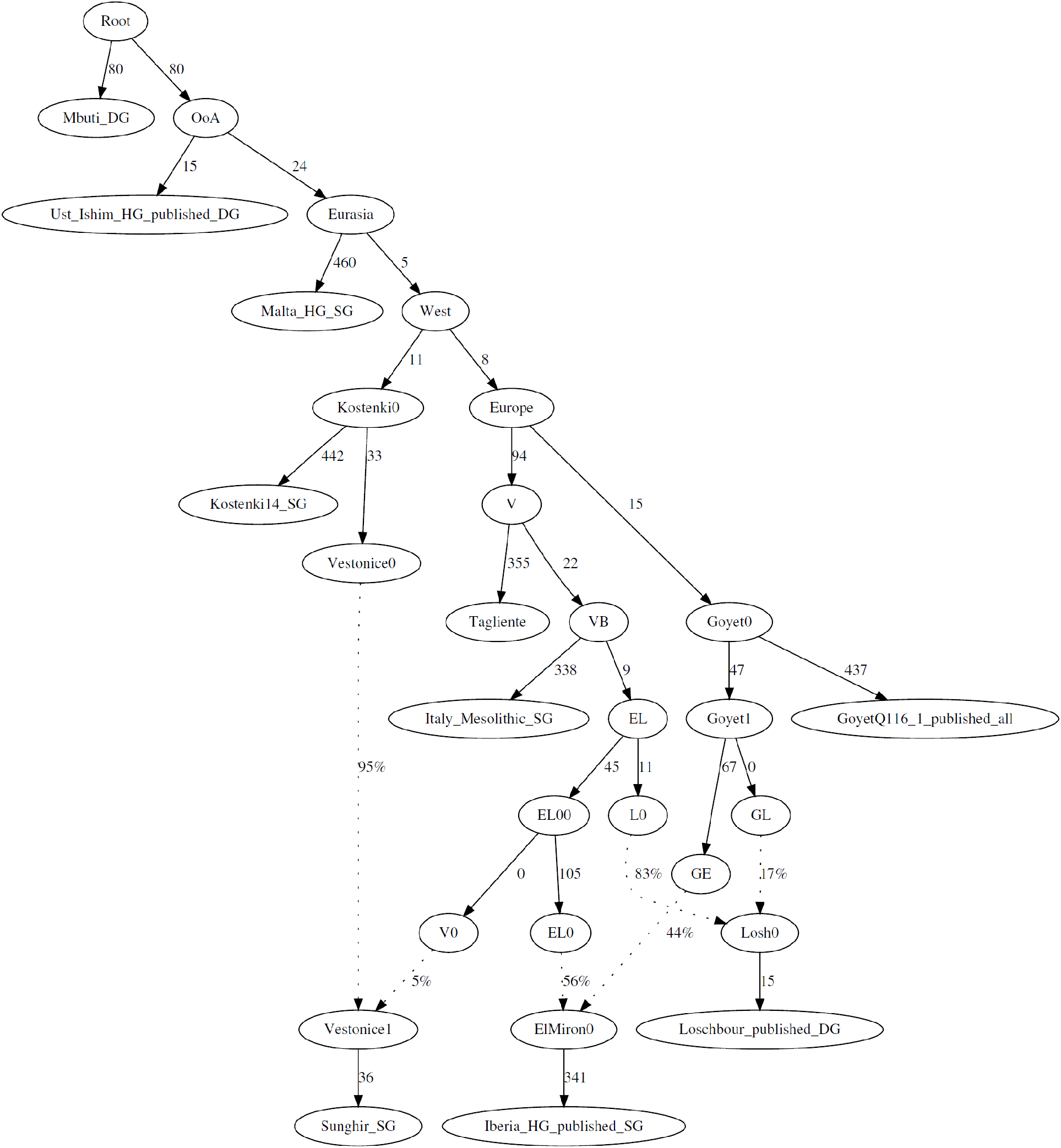
qpGraph using shotgun data. To minimize the bias introduced by using capture and shotgun data within the same analysis, we report also the tree proposed as Extended Data Fig. 4B using, with the exception of Goyet, only shotgun data. Final Score: 34431.549; Degrees of freedom: 15; One f2 outlier: Mal, Sun, Z=2.346; Worst f4: Mbu,Ust,Sun,Goy, Z=3.541.

**Extended Data Fig.6.**
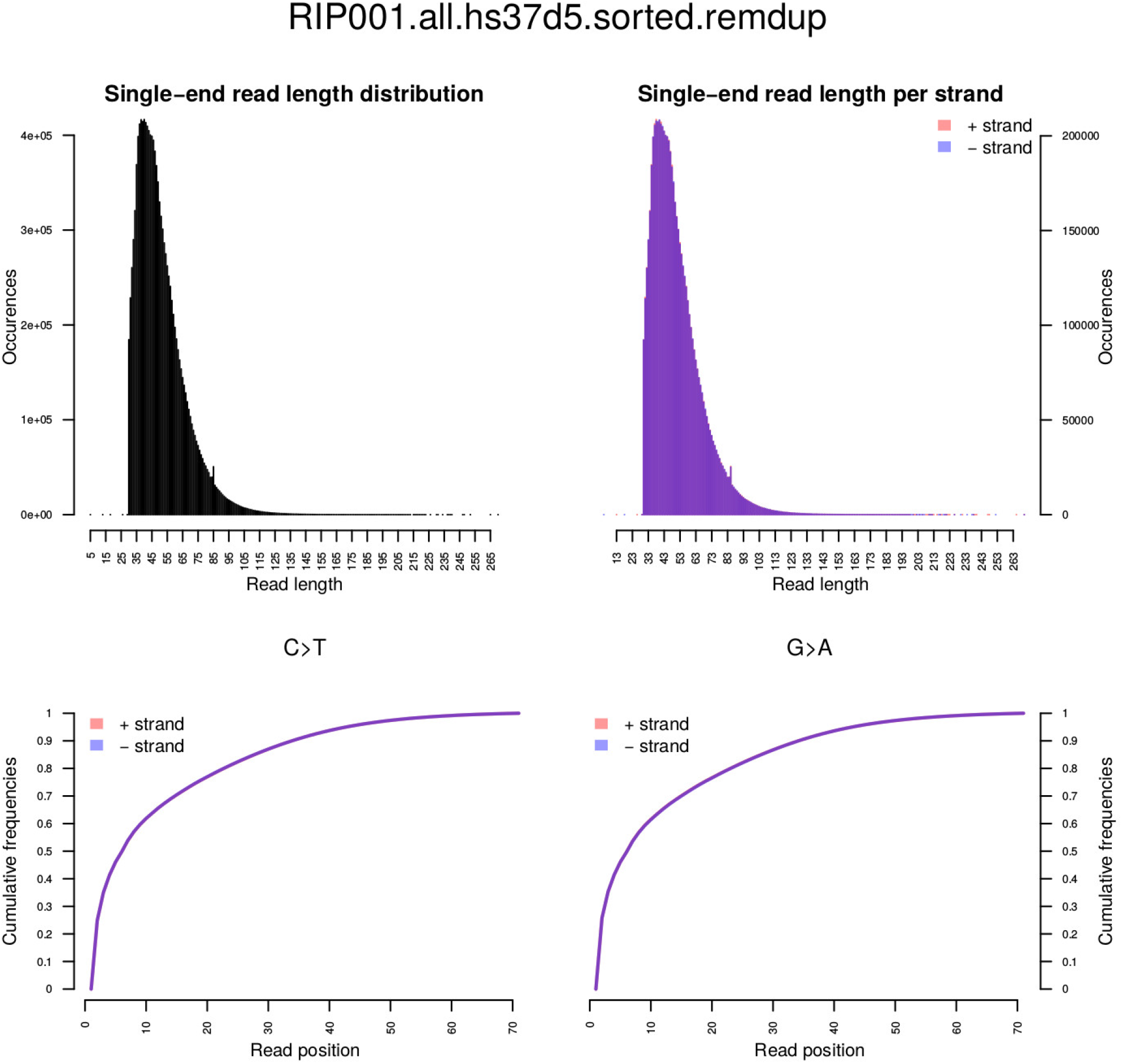
Sequencing read length and substitution rate for Tagliente2 whole genome sequence

## Notes

### Competing Interest Statement

The authors have declared no competing interest.

