## Supplementary Information for "Early Alpine occupation backdates westward human migration in Late Glacial Europe"

**Supplementary Information section 1**
**Palaeoclimate, palaeoenvironmental and bio-cultural changes between 30 and 11 ka**

The period ranging between the Last Glacial Maximum (LGM) and the onset of the Holocene (ca. 30-11 ka cal BP) experienced large-scale climate changes that produced distinctive local and regional ecological and bio-cultural responses (Fig. 1). This time interval is characterised by large migratory events of modern human populations linked to dramatic changes in demography, human behaviour, and the appearance of various material culture complexes (Posth et al., 2016; Fig. 1). Glaciated Alps represented an effective physiographic barrier for meridional moist advection (Luetscher et al., 2015; Lofverstrom, 2020) supporting tree growth and boreal forests in the southern alpine foreland (Pini et al., 2010).

A phase of major forest contraction occurred between ca. 26 to 21 ka cal BP during the GS-3 stadial (Fig. 1c). At this time, European mountain glaciers expanded (see Hughes et al., 2016; Monegato et al., 2017) and large piedmont glaciers advanced onto the Alpine foreland around 25 ky ago (e.g., Ivy-Ochs et al., 2008; Monegato et al., 2017). A continental shelf emerged as consequence of extreme sea level fall (i.e., ~120m; Pellegrini et al. 2015; Maselli et al 2014) connecting the Balkans to the Western Mediterranean regions with a major role in driving large-scale migratory fluxes of Gravettian-Postgravettian hunter-gatherers.

During the LGM culmination there is at present no direct evidence of megafaunal extinction events (Fig.1c). The extinction of Cave bears (Terlato et al., 2018;), is one of the most relevant issues of the Late Quaternary (Cooper et al., 2015) whose main causes are human hunting and interference (Munzel et al., 2011; Romandini and Nannini, 2012; Wojtal et al., 2015; Fortes et al., 2016) and climate change (Pacher and Stuart 2009; Stuart and Lister 2007, Barnosky et al., 2004; Lorenzen et al., 2011; Stuart 2015; Gretzinger et al., 2019). On the other hand, major megafaunal (mainly steppe taxa) extinction events rather appear to be associated with warming events towards the end of the Pleistocene (~14 to 11 ka; Fig.1c).

During the interval ranging between ca. 19-11.7 ka yrs cal BP, the Epigravettian colonization of the Alps, Apennines, and the Dinarids started. After ca. 18 ka, the deglaciation was characterised by dynamic glacial fluctuations (Seguinot et al., 2018; Fig. 1a) through the two combined processes of down melting (in altitudes) and ice-retreat controlled by slope-damming processes along the valley floors. Pine-birch groves and open larch stands established in the South-Alpine foreland, while open woodland of spruce, pine and larch with a juniper understorey colonized ice-free areas in the eastern Pre-Alps (Ravazzi et al. 2014; Vescovi et al., 2007). During this period the first human occurrence at the foot of the Pre-Alps is documented at Riparo Tagliente (Fontana et al., 2009).

This phase was followed by an abrupt shift to warmer conditions at ca. 14.7 ka cal BP (onset of Bølling–Allerød interstadial period, GI-1e in the Greenland ice record) (Fig. 1c), which promoted the displacement of alpine habitats to higher altitudes and an increase in woodland density with dominance of pine, larch, spruce and birch (Vescovi et al., 2007). The second part of this Interstadial was marked by the expansion of mixed oak forests in the Po plain and submountain belt. Such favourable climatic conditions promoted the increase in the number of Epigravettian sites and also a gradual colonization of high altitudes (i.e., above 1000 m a.s.l.). Renewed cold conditions occurred at the beginning of the Younger Dryas (YD, also regarded as event GS 1 in Greenland ice cores; ca. 12.9 -11.7 ka cal BP; e.g., Lowe et al., 2008) (Fig. 1c). It certainly had strong effects on stands of thermophilous trees formerly expanded in the forelands of the Southern Alps. During the second half of this event, a renewed glacial activity is recorded in the high valleys and extends well into the early Holocene (Ivy-Ochs et al., 2008; Moran et al., 2016). This oscillation precedes the final transition to Holocene interglacial conditions at 11.7 ka cal BP. Palaeoecological data indicate a rapid transformation of forests composition in the valley floors and lower slopes. Here, mixed forest with thermophilous trees such as *Quercus*, *Ulmus,* and *Tilia* expanded (Vescovi et al., 2007). Due to the increase of temperature, the timberline rapidly reached an altitude of about 2100 m a.s.l. within few centuries (Oeggl and Wahlmüller 1994). During the

Mesolithic, an intense human colonization of the highlands as well as the valley floors and the middle high landscapes of the Alps and the Italian Prealps is documented (Fontana et al., 2011). In this context, larger residential sites surrounded by more ephemeral sites could be tied together in seasonal vertical mobility along the treeline belt (Fontana and Visentin, 2016).

### Supplementary Information section 2
### Archaeological and ecological context of Tagliente2

Riparo Tagliente (Stallavena di Grezzana, Verona) is located on the left slope of Valpantena, one of the main valley bottoms of the pre-alpine massif of Monti Lessini. The site lies under a rock-shelter and was discovered in 1958 by Mr. Francesco Tagliente. It opens at the base of Monte Tregnago under a bank of oolitic limestones at an altitude of 250 m a.s.l. Archaeological investigations were carried out by the Museo Civico di Storia Naturale of Verona from 1962 to 1964 and resumed by the University of Ferrara since 1967. The site preserves a stratigraphic series which reaches a thickness of over 4.50 m in the outer area of the rock-shelter. Such sequence is formed by two main deposits separated by river erosion: the lower deposit contains evidence of Mousterian and Aurignacian occupations and the upper one is characterized by rich record related to several intense Late Epigravettian settlement occurrences (Bartolomei et al. 1982). According to radiocarbon dates which range from 17,219-16,687 cal BP (layer 13 alpha) to 14,572–13,430 cal BP 14535-13472 (levels 10–8), the Epigravettian series is one of the most complete in northern Italy spanning from the first part of the Lateglacial to the the Bølling–Allerød interstadial (Fontana et al. 2009, 2018).

The hemimandible was found in 1963 during the first excavation campaigns in the site within disturbed sediments located immediately outside the shelter (Corrain 1966). According to excavators such sediments could come from the inner area of the shelter and have been removed during historical excavations in the uppermost deposits which had led to destruction of part of the prehistoric stratigraphic sequence and dumping of sediments outside the shelter entrance. An *in situ* burial was found ten years later (1973) at the southern edge of the sheltered area (Bartolomei et al 1974). The burial had been partially destroyed by the same historical excavations. Nonetheless the lowermost part of the skeleton - from the pelvis to the feet - was well preserved in the grave while only a few ribs and vertebrae, the distal fragments of the radius and ulna and some phalanxes were collected on the trampling floor of the artificial chamber created by historical excavations. The skeleton was contained in a 60 cm deep and 60 cm wide pit with a concave section. It had been laid in a supine position with outstretched arms. The original composition of the grave goods assemblage is unknown due to the incompleteness of the burial but a limestone pebble covered with traces of ochre was found between the feet and a fragment of bovid horn near the right femur. The intentional deposition of a pierced *Cyclope* collected near the left knee is uncertain. The legs were covered with some limestones blocks of different dimensions. A large stone located on the femurs was characterized by engravings of a lion and the horn of an auroch (Bartolomei et al 1974). The skeleton belonged to a young adult male aged 20–29 years and about 163 cm tall (Corrain 1977). It was directly dated from a bone fragment of the right femur to 13,190 ± 90 BP (OxA-10672, 16,634–15,286 cal. BP) (Hedges et al. 1993). Such chronology is close to the one obtained on the hemi-mandible although the two dating do not overlap and it is not possible to affirm that the mandible belongs to the buried individual. In both cases the two specimens belong to adult males. Carbon and nitrogen stable isotope analysis performed on the bone collagen from the partial skeleton (from a rib) and 11 faunal remains from layers consistent with the burial deposition, suggests that the human individual had a terrestrial diet integrated by consumption of aquatic resources (Gazzoni et al. 2013).

Radiocarbon dating of the hemimandible, of the partial burial and the stratigraphic position of the burial pit, which intersects the Mousterian deposits, show consistency with the first phase of Late Epigravettian occupation in the site. At Riparo Tagliente, layers referred to this period are better known in the northern sector where they have been extensively explored in the area protected by the rock-shelter over a surface of around 20 s.q.m. Radiometric dating available for these layers span from 17,219-16,687 cal. BP (layer 13 alpha) to 16438-15941 years cal. BP attesting for the most ancient human presence in the Alpine foothills after the onset of deglaciation (Fontana et al. 2018). Considering the number of determined faunal remains a dominance of species adapted to open environments, such as the ibex and the marmot, is attested in these layers. Further species are represented by red deer (*Cervus elaphus*), roe deer (*Capreolus capreolus*), bison/aurochs (*Bos/Bison*), wild boar (*Sus scrofa*), elk (*Alces sp*.), brown bear (*Ursus arctos*) and chamois (*Rupicapra rupicapra*). Several specimens are characterized by butchering marks and fish and bird remains are also documented, although no specific analysis has been undertaken so far. The analysis of seasonality shows an occupation spanning from the beginning of the spring season to the end of autumn (Fontana et al. 2009, 2018; Gazzoni et al 2013; Rocci Ris 2006). The archeological record testifies for an intense occupation of the site during this period. This record consists of dwelling structures, several fireplaces, and some thick soils rich in lithic assemblages, faunal and ochre remains, along with some osseous tools and some beads obtained from marine shells and red deer canines (Bietti et al. 2004, Cavallo et al. 2017a, 2017b, Fontana et al 2015, Peretto et al. 2004). An emphasis on processing of the rich lithic, mineral and biological resources offered by the Lessini area is recorded. The variety and abundance of finds indicate an excellent knowledge of the territory surrounding the site, which was intensively exploited at least from the valley-bottom to the top of the plateau (Bertola et al. 2007; Fontana et al. 2009). Such occupations occurred on a seasonal basis, especially during the period of the year between early spring and late autumn. Despite this rich record the absence of evidence referring to the time span between 17 and 16 kyrs cal. BP all over the north Italian peninsula does not allow to support any hypothesis on the annual range of mobility of these groups. Nonetheless, the presence of few artefacts and cores manufactured on cherts from the Northern Adriatic Apennines (Umbria-Marche basin) among the wide quantity of items and discarded elements obtained on the local high quality siliceous rocks suggests the persistence of contacts with this area until at least this age (Bertola et al. 2018). Long distance mobility and/or contacts are also supported by marine shells beads from this layers amounting to some hundreds (Fontana et al. 2009)

### Supplementary Information section 3
### Removal of Cementoma

#### *3.1Equipment*

- 24 volt dental micro-motor
- 24 volt power supply with a reduction possibility to 18 and 12 volts
- Single cell charger
- 5x dental red ring contra angle handpiece
- 1/1  dental blue ring contra angle handpiece
- Dental straight handpiece
- Slotted diamond bur 25mm long and 0.5mm tip diameter
- Tungsten carbide bur 31mm long and 0.8mm tip diameter
- Tungsten carbide rosette bur 31mm long and 0.8mm diameter
- Diamond disc for ceramic 0.35mm thick and 20mm diameter

- Blade n°12 surgical scalpel
- Buck 5/6 periodontal scalpel
- The rotation speed of the 1/1 dental blue ring contra angle handpiece is 40.000 rpm, i.e. the motor speed
- The rotation speed of the 5x dental red ring multiplier contra angle handpiece is 200.000 rpm. This speed is reduced to about 150.000 rpm when switching to 18 volts, and to 100.000 rpm when switching to 12 volts
- The rotation speed of the 1/1 dental straight handpiece is 35.000 rpm

#### *3.2 Procedure*

The procedure consisted of creating an area around the exhibit in which a possible introduction of external biological material linked to the extraction procedure was minimized; previously, the exhibit had already been widely contaminated and even likely protected with a vinyl glue thin layer. Each operator wore sterile gown, mask, cap and sterile gloves. All the instruments used, including the handpieces, were sterile; the micro-motors have been sanitized and protected by disposable antiseptic sheaths. The exhibit was placed on a sterile sheet. The procedure was carried out entirely by working with the aid of a Zeiss 6x binocular microscope.

The aim of the project was the extraction of a bone piece containing a pathological alteration (a probable cementoma) located under the buccal vestibulum, slightly distal to half the root height of the lower left second premolar. The other priority of the procedure was the need to minimize the corruption of the exhibit by reducing the operational invasiveness, trying to keep the lingual surface intact, which is the side displayed to museum visitors. Therefore, the difficulty was the extraction of a bone piece limited to the vestibular surface including the lesion and the root portion connected to it without damaging or removing the bone portion and the dental crown above it, also avoiding damaging areas visible to the public.

After a careful tomographic examinations analysis of the exhibit to fully identify the spatial location of the lesion and its relationship with the surrounding bone and dental structures, we moved on to the extraction of the bone piece. The operational phase began with the delimitation of the bone piece by cutting the cortex with a 20mm diameter and 0.35mm thickness diamond disc mounted on a red ring contra angle handpiece and micro-motor set at 18 volts (150.000 rpm). We pick this thickness because a larger one (0.5) would have been too destructive for the exhibit, while a smaller one (0.2) would have made the bone piece subsequent mobilization more difficult. The slag and debris produced in this surface cutting phase were collected in a sterile tube and labeled as “contaminated external slag”. Once the area of about 1cm² was delimited, the grooves were deepened with a 25mm long and 0.5mm diameter conical slotted diamond bur up to a depth of about two thirds of the jaw thickness corresponding to the cortical-spongy bone limit. The pressure exerted with the bur has always been gentle in order to avoid overheating of the bone matrix and loss of diamond crystals. Exploiting the free alveolus of the lower left first molar, as this had been used for previous studies and therefore made removable, it was possible to perform a cut parallel to the lingual surface, orthogonal to the previous perimetral cuts, using a 31mm long and 0.8mm diameter tungsten carbide bur. The slag recovered at this stage has been collected in a sterile tube, and since it had not been contaminated labeled as “useful for microbiological examination”. Given the impossibility to proceed entirely with the diamond tip due to the risk of excessively corrupting the bone matrix, the cuts were defined with a blade n°12 surgical scalpel and a Buck 5/6 periodontal scalpel until complete detachment of the bone piece.

During some separation stages of the internal trabeculation, especially in the periradicular areas, the scalpel blade was gently moved with the aid of small percussions on the handle back, in order to get over the localized resistances that a continuous pressure could not have overcome without significantly increasing strength and dangerously decreasing control. Once all the release incisions were carried out, the bone piece was easily mobilized and extracted with tweezer.

The lesion was therefore exposed by bone matrix abrasion with a 0.8mm tungsten carbide rosette bur mounted on a straight handpiece with a 35.000 rpm micro-motor. This progressive controlled abrasion allowed us to reach the exposure of a lesion portion large enough to be easily accessible for histological and DNA examination.

The entire procedure was documented with a Nikon camera with a 105mm Micro-Nikkor lens equipped with annular flash.

### Supplementary Information section 4
### Genetic analysis of cementoma

We generated 0.28x coverage genome and searched for putatively pathogenic variants in candidate genes (ALPL, ZNF687, CDC73, H3F3A, FOS,H3F3B, FOSB) known to be involved in in similar pathological conditions to  Focal Osseous Dysplasia insurgence or used to perform differential diagnoses (Henthorn et al. 1992; Divisato et al. 2016; Carpten et al. 2002; Behjati et al. 2013; Fittall et al. 2018 ), and found a number of non-reference SNPs which, given the low coverage and ancient DNA degradation are reported here with no further interpretation (Supplementary Table 4).

### Supplementary Information section 5
### Radiocarbon dating of Tagliente1 and Tagliente2

The age of Tagliente2 is 16980-16510years cal BP (95.4% probability). Comparing this result to the one from Tagliente1 (16130-15560cal BP at 95.4% probability; Gazzoni et al. 2013) it appears that the two calibrated radiocarbon ages differ beyond the 2σ level; therefore, we suggest that Tagliente1 and Tagliente2 specimens likely belonged to different individuals. Likewise, stable isotopes of collagen indicate diverse dietary inputs for Tagliente1 (Phoca-Cosmetatou et al. 2005) and 2, corroborating the hypothesis of a different origin for these samples.

**Supplementary Information - References**


Barnosky, A.D., Koch, P.L., Feranec, R.S., Wing, S.L., Shabel, A.B. 2004. Assessing the causes of late Pleistocene extinctions on the continents. Science. 306:70–75.

Bartolomei G, Broglio A, Guerreschi A et al (1974) Una sepoltura epigravettiana nel deposito pleistocenico del Riparo Tagliente in Valpantena (Verona). Rivista di Scienze Preistoriche 29:101–152

Behjati S, Tarpey PS, Presneau N, Scheipl S, Pillay N, Van Loo P, Wedge DC, Cooke SL, Gundem G, Davies H, Nik-Zainal S, Martin S, McLaren S, Goodie V, Robinson B, Butler A, Teague JW, Halai D, Khatri B, Myklebost O, Baumhoer D, Jundt G, Hamoudi R, Tirabosco R, Amary MF, Futreal PA, Stratton MR, Campbell PJ, Flanagan AM. Distinct H3F3A and H3F3B driver mutations define chondroblastoma and giant cell tumor of bone. Nat Genet. 2013, 45:1479-82.

Bertola S, Broglio A, Cassoli PF, et al (2007) L'Epigravettiano recente nell'area prealpina e alpina orientale. In: Martini F (ed) L'Italia tra 15.000 e 10.000 anni fa. Cosmopolitismo e regionalità nel Tardoglaciale. Millenni, Studi di Archeologia Preistoria. Museo Fiorentino di Preistoria "Paolo Graziosi", Firenze, 5, pp 39–94.

Bietti, A., Boschian, G., Mirocle Crisci, G., Danese, E., De Francesco, A. M., Dini, M., Fontana, F., Giampietri, A., Grifoni, R., Guerreschi, A., Liagre, J., Negrino, F., Radi, G., Tozzi, C. and Tykot, R. 2004. Inorganic raw materials economy and provenance of chipped industry in some stone age sites of Northern and Central Italy. Collegium Anthropologicum, 28,1, pp. 41-54

Brock, F., Bronk Ramsey, C., Higham, Quality assurance of ultrafiltered bone dating. Radiocarbon. 49, 187–192 (2007)

Brown, T.A., Nelson, D.E., Vogel, J.S., Southon, J.R. Improved Collagen Extraction by modified Longin method. Radiocarbon. 30, 171-177 (1988)

Carpten JD, Robbins CM, Villablanca A, Forsberg L, Presciuttini S, Bailey-Wilson J, Simonds WF, Gillanders EM, Kennedy AM, Chen JD, Agarwal SK, Sood R, Jones MP, Moses TY, Haven C, Petillo D, Leotlela PD, Harding B, Cameron D, Pannett AA, Höög A, Heath H 3rd, James-Newton LA, Robinson B, Zarbo RJ, Cavaco BM, Wassif W, Perrier ND, Rosen IB, Kristoffersson U, Turnpenny PD, Farnebo LO, Besser GM, Jackson CE, Morreau H, Trent JM, Thakker RV, Marx SJ, Teh BT, Larsson C, Hobbs MR. HRPT2, encoding parafibromin, is mutated in hyperparathyroidism-jaw tumor syndrome. Nature Genet. 2002, 32: 676-680, 2002.

Cavallo, G., Fontana, F., Gonzato, F., Guerreschi, A., Riccardi, M.P., Sardelli, G. and Zorzin, R. 2017a. Sourcing and processing of ochre during the late upper Palaeolithic at Tagliente rock-shelter (NE Italy) based on conventional X-ray powder diffraction analysis. Archaeological and Anthropological Sciences, 9 (5), pp. 763-775.

Cavallo, G., Fontana, F., Gonzato, F., Peresani, M., Riccardi, M.P and Zorzin, R. 2017b. Textural, microstructural, and compositional characteristics of Fe-based geomaterials and Upper Palaeolithic ochre in the Lessini Mountains, NE Italy: Implications for provenance studies" for publication in Geoarchaeology. Geoarcheology, 32 (4), pp. 437-455.

Cooper, A., Turney, C., Hughen, K.A., Barry W. Brook B.W., McDonald, H.G., Bradshaw C.J.A., 2015. Abrupt warming events drove Late Pleistocene Holarctic megafaunal turnover. Science; Vol. 349, Issue 6248, pp. 602-606. DOI: 10.1126/science.aac4315

Corrain C (1977) I resti scheletrici umani della sepoltura epigravettianadel Riparo Tagliente in Valpantena (Verona). Bollettino del Museo Civico di Storia Naturale di Verona 4:35–79

Divisato G, Formicola D, Esposito T, Merlotti D, Pazzaglia L, Del Fattore A, Siris E, Orcel P, Brown JP, Nuti R, Strazzullo P, Benassi MS, Cancela ML, Michou L, Rendina D, Gennari L, Gianfrancesco F. ZNF687 Mutations in Severe Paget Disease of Bone Associated with Giant Cell Tumor. Am J Hum Genet. 2016, 98:275-86.

Fittall MW, Mifsud W, Pillay N, Ye H, Strobl AC, Verfaillie A, Demeulemeester J, Zhang L, Berisha F, Tarabichi M, Young MD, Miranda E, Tarpey PS, Tirabosco R, Amary F, Grigoriadis AE, Stratton MR, Van Loo P, Antonescu CR, Campbell PJ, Flanagan AM, Behjati S. Recurrent rearrangements of FOS and FOSB define osteoblastoma. Nat Commun. 2018, 9:2150.

Fontana, F., Cilli, C., Cremona, M.G., Giacobini, G., Gurioli, F., Liagre, J., …..,Guerreschi, A. (2009). Recent data on the Late Epigravettian occupation at Riparo Tagliente, Monti Lessini (Grezzana, Verona): a multidisciplinary perspective. Preistoria Alpina, 44, 51-59 (ISSN 0393-0157).

Fontana, F., Guerreschi, A., & Peresani, M. (2011). The visible landscape: Inferring Mesolithic settlement dynamics from multifaceted evidence in the south-eastern Alps. In Hidden Landscapes of Mediterranean Europe: Cultural and Methodological Biases in Pre-and Protohistoric Landscape Studies. Proceedings of the International Meeting (Siena 2007), Oxford, BAR International Series (Vol. 2320, pp. 71-81).

Fontana, F., Guerreschi, A., Bertola, S., Cremona, M.G., Cavulli, F., Falceri, L., Gajardo, A., Montoya, C., Ndyaie, M. and Visentin, D. 2015. I livelli più antichi della serie epigravettiana “interna” di Riparo Tagliente: sfruttamento delle risorse litiche e sistemi tecnici. In: Leonardi, G. and Tinè, V. eds. Atti della XLVIII Riunione Scientifica dell’Istituto Italiano di Preistoria e Protostoria, Preistoria e Protostoria del Veneto, Padova 5-9 novembre 2013, Istituto Italiano di Preistoria e Protostoria, pp. 43-52.

Fontana, F., & Visentin, D. (2016). Between the Venetian Alps and the Emilian Apennines (Northern Italy): Highland vs. lowland occupation in the early Mesolithic. Quaternary international, 423, 266-278.

Fortes, G.G., Grandal-d’Anglade, A., Kolbe, B., Fernandes, D., Meleg, I.N., Garcia-Vazquez, A., Pinto-Llona, A.C., Constantin, S., de Torres, T.J., 2016. Ancient DNA reveals differences in behaviour and sociality between brown bears and extinct cave bears. Mol Ecol. 25:4907–4918.

Gazzoni, V., et al. (2013). Late Upper Palaeolithic human diet: first stable isotope evidence from Riparo Tagliente (Verona, Italy). Bull. Mém. Soc. Anthropol. 25, 103-117

Gretzinger J., Molak M., Reiter E., Pfrengle S., Urban C., Neukamm J., Blant M., Conard N.J., Cupillard C.,Dimitrijević V., Drucker D.G., Hofman-Kamiń ska E., Kowalczyk R., Krajcarz M.T., Krajcarz M., Münzel S.C., Peresani M., Romandini M., Rufí I., Soler J., Terlato G., Krause J., Bocherens H., Schuenemann V.J., 2019. Large-scale mitogenomic analysis of the phylogeography of the Late Pleistocene cave bear. Sci Rep 9, 10700 (2019). https://doi.org/10.1038/s41598-019-47073-z

Hedges R, Housley RA, Bronk Ramsey C, et al (1993) Radiocarbon dates from the Oxford AMS system: Archaeometry Datelist 16. Archaeometry 35/1:147–67

Henthorn, P. S., Raducha, M., Fedde, K. N., Lafferty, M. A., Whyte, M. P. Different missense mutations at the tissue-nonspecific alkaline phosphatase gene locus in autosomal recessively inherited forms of mild and severe hypophosphatasia. Proc. Nat. Acad. Sci.1992, 89: 9924-9928.

Higham, F. G., Jacobi, R. M., Bronk Ramsey, C. AMS radiocarbon dating of ancient bone using ultrafiltration. Radiocarbon. 48, 179-195 (2006)

Ivy Ochs, S., Kerschner, H., Reuther, A., Preusser, F., Heine, K., Maisch, M., ... & Schlüchter, C. (2008). Chronology of the last glacial cycle in the European Alps. Journal of Quaternary Science: Published for the Quaternary Research Association, 23(6 7), 559-573.

Korlević, P., Talamo, S., Meyer, M. A combined method for DNA analysis and radiocarbon dating from a single sample. Sci. Rep. 8, 4127 (2018)

van Klinken, G. J. Bone Collagen Quality Indicators for Palaeodietary and Radiocarbon Measurements. J Archaeol. Sci. 26, 687-695 (1999)

Kromer, B., Lindauer, S., Synal, H.-A., Wacker, L. MAMS – A new AMS facility at the Curt-Engelhorn-Centre for Achaeometry, Mannheim, Germany. Nucl. Instrum. Meth. B. 294, 11-13 (2013)

Longin, R. New method of collagen extraction for radiocarbon dating. Nature. 230, 241-242 (1971)

Phoca-Cosmetatou, N., Landscape use in Northeast Italy during the Upper Palaeolithic. Preistoria Alpina. 41, 23-49 (2005a)

Lorenzen, E.D., Nogues-Bravo, D., Orlando, L., Weinstock, J., Binladen, J., Marske, K.A., Ugan, A., Borregaard, M.K., Gilbert, M.T.P., Nielsen, R., et al. 2011. Species-specific responses of Late Quaternary megafauna to climate and humans. Nature. 479:359–364.

Lofverstrom, M. (2020). A dynamic link between high-intensity precipitation events in southwestern North America and Europe at the Last Glacial Maximum. *Earth and Planetary Science Letters, 534*, 116081.

Lowe, J. J., Rasmussen, S. O., Björck, S., Hoek, W. Z., Steffensen, J. P., Walker, M. J., ... & Intimate Group. (2008). Synchronisation of palaeoenvironmental events in the North Atlantic region during the Last Termination: a revised protocol recommended by the INTIMATE group. Quaternary Science Reviews, 27(1-2), 6-17.

Luetscher, M., Boch, R., Sodemann, H., Spötl, C., Cheng, H., Edwards, R. L., ... & Müller, W. (2015). North Atlantic storm track changes during the Last Glacial Maximum recorded by Alpine speleothems. Nature Communications, 6(1), 1-6.

Maselli, V., Trincardi, F., Asioli, A., Ceregato, A., Rizzetto, F., & Taviani, M. (2014). Delta growth and river valleys: the influence of climate and sea level changes on the South Adriatic shelf (Mediterranean Sea). Quaternary Science Reviews, 99, 146-163.

Monegato, G., Scardia, G., Hajdas, I., Rizzini, F., & Piccin, A. (2017). The Alpine LGM in the boreal ice-sheets game. Scientific reports, 7(1), 1-8.

Moran, A. P., Ivy Ochs, S., Schuh, M., Christl, M., & Kerschner, H. (2016). Evidence of central Alpine glacier advances during the Younger Dryas–early Holocene transition period. Boreas, 45(3), 398-410.

Munzel, S.C., Stiller, M., Hofreiter, M., Mittnik, A., Conard, N.J., Bocherens, H. 2011. Pleistocene bears in the Swabian Jura (Germany): genetic replacement, ecological displacement, extinctions and survival. Quat. Int. 245:1–13.

Oeggl, K., & Wahlmüller, N. (1994). Vegetation and climate history of a high alpine mesolithic camp site in the Eastern Alps. Museo Tridentino di Scienze Naturali.

Pacher, M., Stuart, A.J., 2009. Extinction chronology and palaeobiology of the cave bear (Ursus spelaeus). Boreas. 38:189–206.

Pellegrini, C., Maselli, V., Cattaneo, A., Piva, A., Ceregato, A., & Trincardi, F. (2015). Anatomy of a compound delta from the post-glacial transgressive record in the Adriatic Sea. Marine Geology, 362, 43-59.

Peretto, C., Biagi, P., Boschian, G., Broglio, A., De Stefani, M., Fasani, L., Fontana, F., Grifoni, R., Guerreschi, A., Iacopini, A., Minelli, A., Pala, R., Peresani, M., Radi, G., Ronchitelli, A., Sarti, L., Thun Hohenstein, U. and Tozzi C. 2004. Living-floors and structures from the lower Palaeolithic to the Bronze age in Italy. Collegium Anthropologicum 28 (1) pp. 63-88.

Phoca-Cosmetatou, N. Bone weathering and food procurement strategies: assessing the reliability of our behavioural inferences. Biosphere to Lithosphere: New Studies in Vertebrate Taphonomy. (ed. O'Connor, T.P.) 135-145. (Oxbow Books, 2005b)

Pini, R., Ravazzi, C., & Reimer, P. J. (2010). The vegetation and climate history of the last glacial cycle in a new pollen record from Lake Fimon (southern Alpine foreland, N-Italy). *Quaternary Science Reviews*, *29*(23-24), 3115-3137.

Posth, C., Renaud, G., Mittnik, A., Drucker, D. G., Rougier, H., Cupillard, C., ... & Francken, M. (2016). Pleistocene mitochondrial genomes suggest a single major dispersal of non-Africans and a Late Glacial population turnover in Europe. Current Biology, 26(6), 827-833.

Ravazzi, C., Pini, R., Badino, F., De Amicis, M., Londeix, L., & Reimer, P. J. (2014). The latest LGM culmination of the Garda Glacier (Italian Alps) and the onset of glacial termination. Age of glacial collapse and vegetation chronosequence. Quaternary Science Reviews, 105, 26-47.

Rocci Ris, A. 2006. I macromammiferi di Riparo Tagliente. Archeozoologia e tafonomia dei livelli epigravettiani. PhD thesis, Dipartimento di Anatomia, Farmacologia e medicina Legale, Università degli Studi di Torino.

Seguinot, J., Ivy-Ochs, S., Jouvet, G., Huss, M., Funk, M., & Preusser, F. (2018). Modelling last glacial cycle ice dynamics in the Alps. The Cryosphere, 12(10), 3265-3285.

Stiner, M.C., Kuhn, S.L., Weiner, S., Bar-Yosef, O. Differential Burning, Recrystallization and Fragmentation of Archaeological Bone. J. Archaeol. Sci. 22, 223-237 (1995)

Stuart, A.J., 2015. Late Quaternary megafaunal extinctions on the continents: a short review. Geol J. 50:338–363.

Stuart, A.J., Lister, A.M., 2007. Patterns of Late Quaternary megafaunal extinctions in Europe and northern Asia. Cour Forsch-Inst Senckenberg. 259:287–297.

Vescovi, E., Ravazzi, C., Arpenti, E., Finsinger, W., Pini, R., Valsecchi, V., ... & Tinner, W. (2007). Interactions between climate and vegetation during the Lateglacial period as recorded by lake and mire sediment archives in Northern Italy and Southern Switzerland. Quaternary Science Reviews, 26(11-12), 1650-1669.

Wick, L. Early-Holocene reforestation e vegetation change at a lake near the Alpine forest limit: Lago Basso (2250 m asl), Northern Italy. (ed. A. F. Lotter e B. Ammann) 555-563. (Dissertationes Botanicae, 1994)
